# KAT8 compound inhibition inhibits the initial steps of PINK1-dependant mitophagy

**DOI:** 10.1101/2023.08.03.551835

**Authors:** Capucine de Talhouët, Noemi Esteras Gallego, Marc Soutar, Benjamin O’Callaghan, Helene Plun-Favreau

## Abstract

It has recently been shown that *KAT8*, a genome-wide association study (GWAS) candidate risk gene for Parkinson’s Disease, is involved in PINK1/Parkin-dependant mitophagy. The *KAT8* gene encodes a lysine acetyltransferase and represents the catalytically active subunit of the non-specific lethal (NSL) epigenetic remodelling complex. In the current study, we show that contrary to KAT5 inhibition, dual inhibition of KAT5 and KAT8 via the MG149 compound inhibits the initial steps of the PINK1-dependant mitophagy process. More specifically, our study shows that following mitochondrial depolarisation induced by mitochondrial toxins, MG149 treatment inhibits PINK1-dependant mitophagy initiation by impairing PINK1 activation, and subsequent phosphorylation of Parkin and ubiquitin. While this inhibitory effect of MG149 on PINK1-activation is potent, MG149 treatment in the absence of mitochondrial toxins is sufficient to depolarise the mitochondrial membrane, recruit PINK1 and promote partial downstream recruitment of the autophagy receptor p62, leading to an increase in mitochondrial delivery to the lysosomes. Altogether, our study provides additional support for KAT8 as a regulator of mitophagy and autophagy processes.

## Introduction

Like other neurodegenerative diseases, Parkinson’s disease (PD) can either be inherited or stem from more variable sporadic origins. The Mendelian forms of PD have been of major importance for a better understanding of the pathophysiology of the disease. A few of the causative Mendelian PD mutations have been identified in genes encoding proteins which play a key role in the mitophagy process, namely *PINK1* and *PRKN*, respectively encoding a serine/threonine kinase (PTEN-induced kinase 1) and an E3 ubiquitin ligase (Deas, E., *et al*., 2011 ^2^) (Park, J., *et al*, 2006) (Clark, I. E., *et al*., 2006). This identified dysfunctional mitophagy as an important underlying pathomechanism in familial PD.

Mitochondria are essential for a wide range of cellular processes such as ATP production through oxidative phosphorylation, calcium homeostasis, reactive oxygen species (ROS) production, release and consumption, and apoptosis (López-Doménech, G., *et al*., 2021) (Brand, M. D., *et al*., 2013). When heavily damaged, mitochondria can release toxic ROS throughout the cell, promoting oxidative stress and ultimately cell death (Guo, C. Y., *et al*., 2013). As a result, mitochondrial quality control processes are essential, serving to identify, isolate, and remove damaged mitochondria, thus limiting the accumulation of toxic mitochondrial components (Pickles *et al*., 2018).

PINK1/Parkin-dependant mitophagy is one of several types of selective autophagy aimed towards the degradation of heavily toxic mitochondria (Wang, L., *et al*., 2020) (O’Callaghan, B, *et al*., 2023). In cells with a healthy mitochondrial network, PINK1 is maintained at low levels through constitutive and rapid degradation by the proteasome via the N-end rule pathway (Yamano, K., *et al*., R. J., 2013). Upon mitochondrial damage and associated depolarisation of the mitochondrial membrane, PINK1 accumulates at the mitochondrial surface, where it becomes activated through dimerization and autophosphorylation (Rasool, S., *et al*., 2018). Activated PINK1 phosphorylates ubiquitin already present at the outer mitochondrial membrane (OMM) at serine 65 (pUb(Ser65)) (Kazlauskaite, A., *et al*., 2014) (Kane, L. A., *et al*., 2014) (Koyano, K., *et al*., 2014) (Shiba-Fukushima, K., *et al*., 2014). Parkin is then recruited from the cytosol to those pUb(Ser65) sites and is subsequently phosphorylated by PINK1 at serine 65 (Kazlauskaite, A., *et al*., 2014) (Kondapalli, C., *et al*., 2012). PINK1-dependant phosphorylation of Parkin (pParkin) is required for the full activity of the E3 ubiquitin ligase (McWilliams, T. G., *et al*., 2018) (Wang, L., *et al*., 2020). pParkin mediates the ubiquitination of a wide range of OMM proteins, generating more ubiquitinated sites for PINK1 to phosphorylate, creating a phosphorylation / ubiquitination positive feedback loop at the OMM of damaged mitochondria. Mitofusin 2 (MFN2), one of the substrates for endogenous Parkin (Jin, S. M., *et al*., 2012), undergoes ubiquitination and degradation in early stages of mitophagy and is therefore often used as a marker for mitophagy initiation (Wang, L., *et al*., 2020). Autophagy adapters such as p62 can bind to pUb(Ser65) ubiquitin chains, facilitating autophagosome formation around the mitochondria and ultimate fusion to the lysosomes for degradation (Tanida, I., *et al*., 2008).

Recent description of impairments in PINK1-dependant mitophagy associated with loss of function in idiopathic PD GWAS risk gene candidates *KANSL1* and *KAT8* has suggested impairments in PINK1-mitophagy in the more common idiopathic forms of the disease (Soutar, M., *et al*., 2022). Lysine Acetyltransferase-8 (KAT8) serves as the catalytic acetyltransferase subunit of the male-specific lethal (MSL) and the non-specific lethal (NSL) complex (Radzisheuskaya, A., *et al*., 2021), and is a key player in the regulation of nuclear gene transcription (Sheikh, B. N., *et al*., 2019). KAT8 was shown to regulate PINK1-Parkin dependant mitophagy by regulating *PINK1* gene transcript levels, levels of PINK1 protein and subsequent PINK1-dependant phosphorylation of Parkin and ubiquitin (Soutar, M., *et al*., 2022).

The lysine acetylation process involves the exchange of a H^+^on the ε-position NH3+ of lysine for an acetyl group of an acetyl co-enzyme A (acetyl-CoA). This results in neutralization of lysine residues positive charge and in turn protein conformational changes, which may amongst other things impact regulation of enzymatic activity (Farria, A., *et al*., 2015). The possibility to modulate the activity of lysine acetyltransferases (KATs) and lysine deacetylases (KDACs) demonstrates the potential of protein acetylation as a “regulatory switch” for cellular metabolic processes (Farria, A., *et al*., 2015), (Xu, W., *et al*., 2014), (Christensen, D. G., *et al*., 2019). Modulating the inhibition or activation of different KATs and KDACs has been previously used in cancer therapies (Farria, A., *et al*., 2015). This knowledge, combined with the newfound importance of KAT8 in PINK1-dependant mitophagy, prompted us to investigate the impact of small molecule KAT8 inhibitors on PINK1-dependant mitophagy. Of particular interest for this study is MG149, a dual KAT5 and KAT8 inhibitor whose inhibition is competitive towards the acetyl-CoA binding site in both KAT5 and KAT8 (Ghizzoni, M., *et al*., 2012). KAT5 is another lysine acetyltransferase involved in cellular processes including transcriptional regulations and DNA damage repair. Due to the unavailability of a KAT8 specific inhibitor, we used a specific KAT5 inhibitor, NU9056, as a control for MG149’s activity mediated through KAT5 (Wang J., *et al*., 2020).

Here we show that the effect of MG149 on mitophagy is two-fold: 1) it inhibits PINK1-dependant mitophagy initiation by decreasing PINK1 kinase activity and 2) it induces mitochondrial depolarisation and PINK1-dependant mitochondrial delivery to the lysosomes.

## Results

### MG149 treatment reduces PINK1/Parkin-dependant mitophagy initiation

The consequence of MG149 treatment on PINK1-dependant mitophagy initiation was first determined by assessing PINK1-dependant pUb(Ser65) following mitochondrial depolarisation. The effect of MG149 and NU9056 on pUb(Ser65) deposition was tested by immunofluorescence in Flag-Parkin over-expressing (POE) SH-SY5Y neuroblastoma cells. POE SH-SY5Ys were pre-treated for 1, 3, 24, and 48hr with either 10µM NU9056 (targeting KAT5 at IC_50_ of 2µM, 100µM of NU9056 proved toxic for cells) (Li, L., *et al*., 2020) or 100µM MG149 (targeting KAT5 and KAT8 at IC_50_ of 74µM and 47µM respectively) (Wang J., *et al*., 2020), prior to inducing mitophagy with a combination of 1µM oligomycin (an inhibitor of ATP synthase / mitochondrial complex V) and 1µM antimycin (an inhibitor of the mitochondrial complex III) (O/A) for 3hr. O/A-induced mitochondrial depolarisation induced PINK1-mediated pUb(Ser65) (Figure 1A top panel, quantified in Figure 1B), as described previously (Kazlauskaite, A., *et al*., 2014). 1hr and 3hr MG149 pre-treatments decreased O/A-induced pUb(Ser65) (Figure 1A,B), and did so in a dose-dependant manner (Supplementary Figure 1B), whereas NU9056 had no effect at any time point or dose (Figure 1A,B, Supplementary Figure 1A). Notably, 100μM MG149 pre-treatment for 1hr and 3hr led to a small, non-significant increase in pUb(Ser65) deposition in the absence of O/A treatment (Figure 1A bottom left panel, quantified in Figure 1B). This increase in pUb(Ser65) deposition was also observed with MG149 concentrations >40μM (Supplementary Figure 1B). These data suggest that while MG149 pre-treatment decreases PINK1-dependant pUb(Ser65) deposition following O/A treatment, treatment with MG149 alone is sufficient to induce a subtle pUb(Ser65) deposition in the absence of O/A. The inhibitory effect of MG149 on pUb(Ser65) levels was confirmed by immunoblotting of POE SH-SY5Y cells treated for 3hr with 100µM of MG149 prior to 3hr treatment with O/A (Figure 1C,D). The inhibitory effect of MG149 on PINK1/Parkin-dependant mitophagy initiation was further validated using other markers, namely PINK1-dependant phosphorylation of Parkin at serine 65 (pParkin(Ser65)), and ubiquitination of mitofusin 2 (MFN2) (Figure 1C,D). Importantly, the KAT5 inhibitor NU9056 had no effect on any of these mitophagy markers, strongly suggesting that PINK1-dependant mitophagy initiation is modulated by KAT8, as opposed to KAT5, as shown previously (Soutar, M., *et al*., 2022).

**Figure 1.**
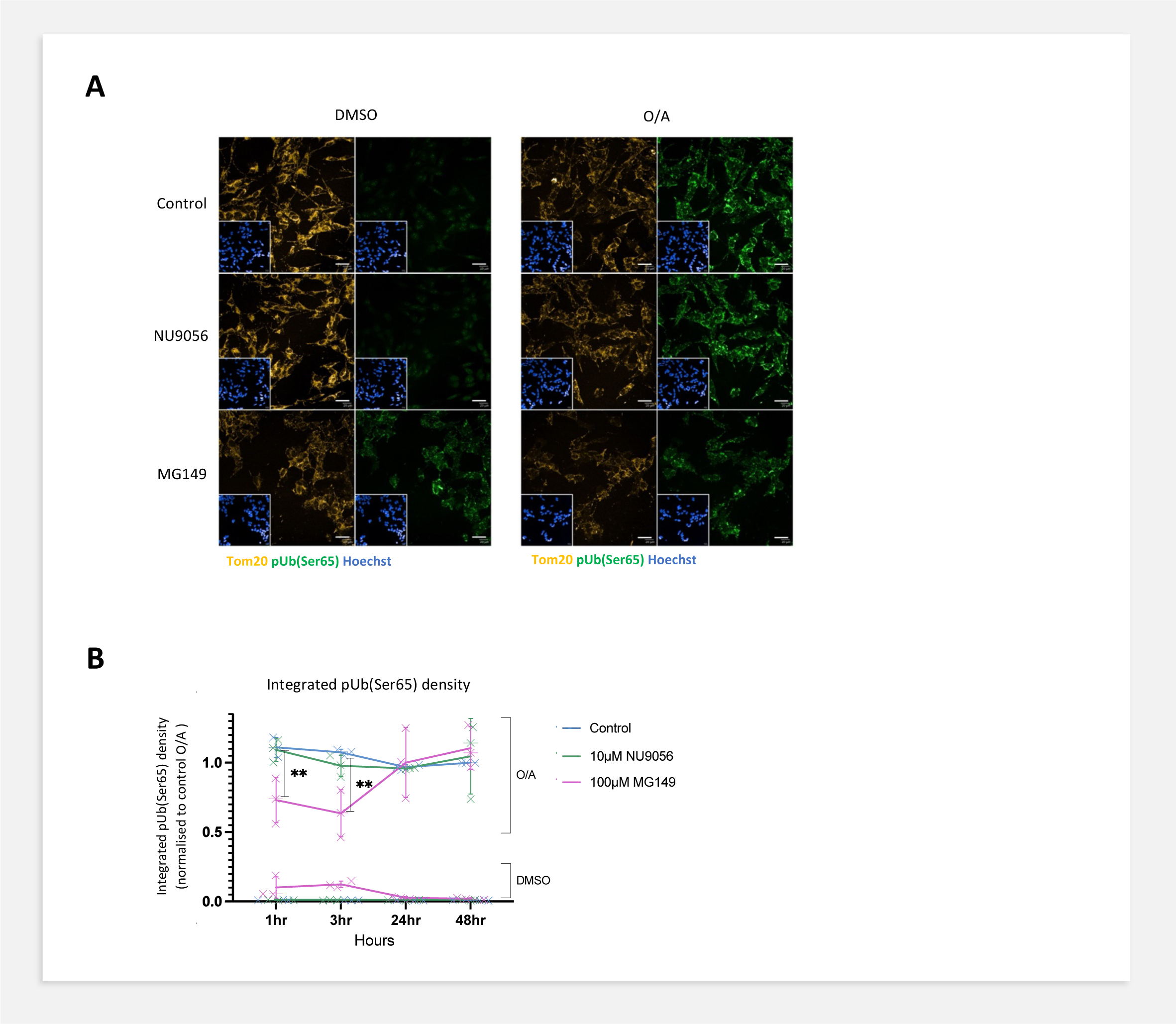
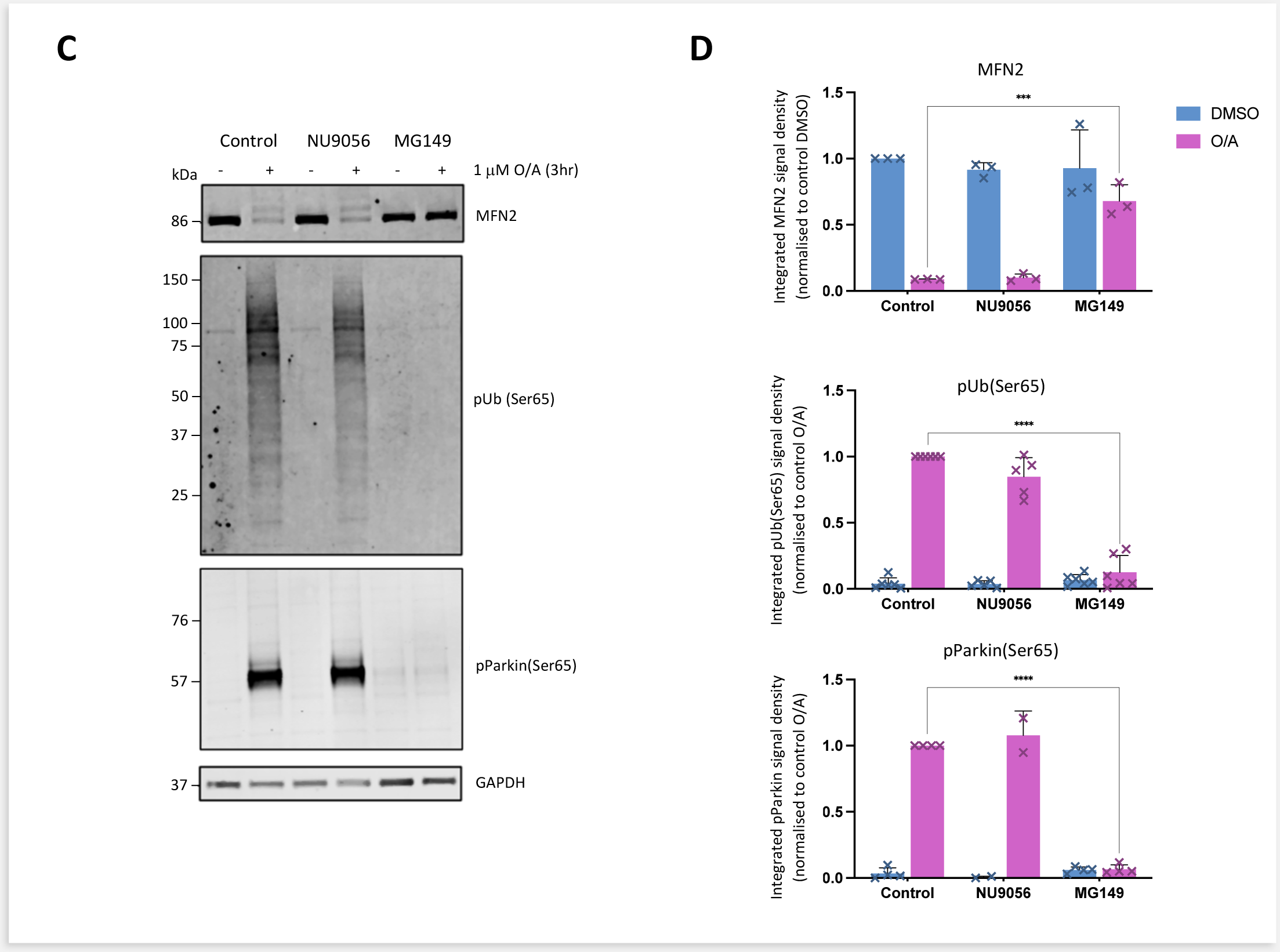
MG149 treatment reduces PINK1/Parkin-dependant mitophagy initiation. **A.** Representative images of POE SH-SY5Y cells pre-treated with DMSO, 10µM NU9056 (KAT5 inhibitor) or 100µM MG149 (KAT5/KAT8 inhibitor) for 1, 3, 24, and 48hr and subsequently treated with DMSO or 1µM O/A for 3hr to induce mitophagy (representative images show a 3hr timepoint). Cells were then immuno-stained anti-Tom20 (yellow) and anti-pUb(Ser65) (green), with Hoechst nuclear counterstain (blue). Scale bar: 20µm. **B.** Quantification of integrated pUb(Ser65) signal density in A, compared to control conditions (n=3, two-way ANOVA with Dunnett’s correction). Data normalized to control O/A (specifically the 48hr DMSO control pre-treatment and 3hr O/A treatment timepoint). **C.** Representative immunoblots of POE SH-SY5Y cells pre-treated with DMSO, 10µM NU9056, 100µM MG149 for 3hr and subsequently treated with DMSO or 1µM O/A for 3hr. Blots were probed for MFN2, pUb(Ser65), pParkin(Ser65) and GAPDH. **D.** Quantification of MFN2, pUb(Ser65), pParkin signal density in C, compared to control conditions (n=3-5, two-way ANOVA with Dunnett’s correction). Data normalized to control O/A and GAPDH used as a loading control. Data shown as mean +/- SD.

### MG149 -treatment does not prevent lysosomal mitochondrial clearance

In order to understand whether the impairments in PINK1-mitophagy initiation observed following MG149 treatment were associated with reductions in delivery of mitochondria to the lysosomes for clearance, we used the pH-sensitive mitochondrial probe mitoKeima, which shifts fluorescence from green to red as mitochondria are delivered to the lysosomes, enabling the measurement of mitochondrial delivery to the lysosomes (Keima mitophagy index) (Katayama, H., *et al*., 2011). POE SH-SY5Ys were simultaneously treated with either 10µM NU9056 or 100µM MG149, and with either DMSO or 1µM O/A, prior to live imaging for 9hr. MG149 did not alter the Keima mitophagy index of O/A-treated cells, suggesting that despite the reduction in PINK1-dependant activation of mitophagy initiation, mitochondrial delivery to the lysosomes was not affected. On the other hand, MG149-treatment led to a significant increase in mitochondrial delivery to the lysosome in the absence of O/A treatment (Figure 2A,B). This increase in mitochondrial delivery was further confirmed using the mitoSRAI mitophagy reporter (Katayama, H., *et al*., 2020). mitoSRAI is a tandem-fluorophore construct targeted to the mitochondrial matrix which consists of the yellow fluorescence protein YPet (sensitive to lysosomal degradation), and TOLLES, a cyan fluorescent protein (insensitive to lysosomal degradation). Tracking the proportion of TOLLES positive mitochondria which have lost Ypet fluorescence enables the measurement of lysosomal delivery of mitochondria (SRAI mitophagy index). Cells were pre-treated for 3hr with either 10µM NU9056 or 100µM MG149, prior to treatment with DMSO or 1µM O/A for 3hr. Similar to the effect observed using mitoKeima, MG149 did not prevent O/A-induced lysosomal delivery of mitochondria while promoting a significant increase in mitochondrial delivery to the lysosome in the absence of O/A treatment (Figure 2C,D).

**Figure 2.**
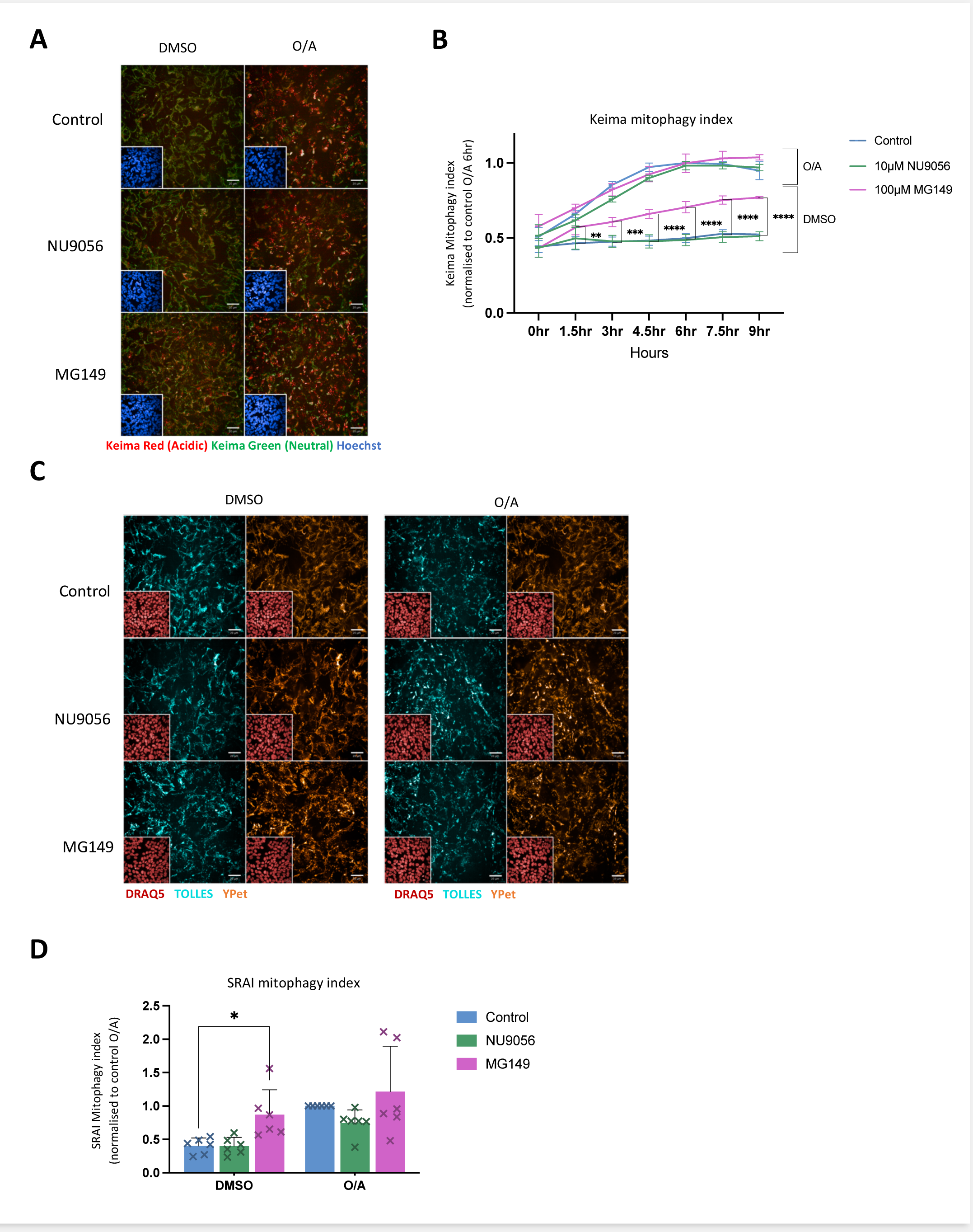
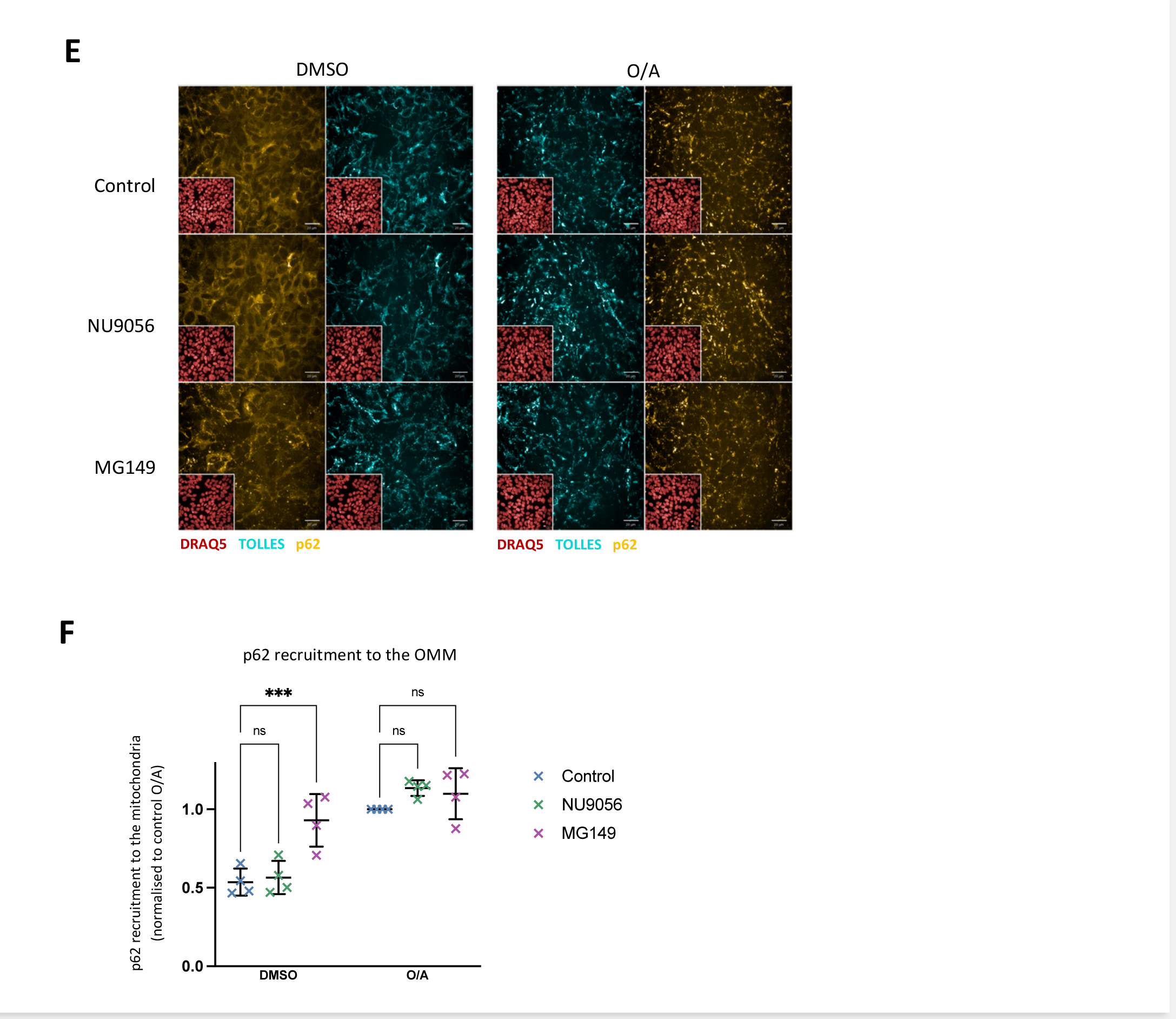
MG149 treatment initiates mitochondrial delivery to the lysosome. **A.** Live time-course imaging of mitoKeima POE SH-SY5Y cells simultaneously treated with DMSO, 10µM NU9056 or 100µM MG149 and with either DMSO or 1µM O/A to induce mitophagy (representative images show a 6hr time-point). The neutral Keima-green signal (green) is shown alongside the acidic lysosomal Keima-red signal (red). Scale bar: 20µm. **B.** Quantification of the Keima mitophagy index in A, calculated as the ratio of the area of lysosomal mitoKeima signal to total mitoKeima signal (sum of lysosomal and neutral mitoKeima signal), compared to control conditions (n=4, two-way ANOVA with Dunnett’s correction). Data normalised to control 6hr O/A. **C.** Representative images of mitoSRAI POE SH-SY5Y cells pre-treated with DMSO, 10µM NU9056 or 100µM MG149 for 3hr and subsequently treated with DMSO or 1µM O/A for 3hr. Cell nuclei were counterstained with DRAQ5 (red). The TOLLES fluorophore (cyan) shows total mitochondria while the Ypet fluorophore (orange) shows the non-lysosomal mitochondria. Scale bar: 20µm. **D.** Quantification of the SRAI mitophagy index in C, calculated as the ratio of the lysosomal mitochondria (TOLLES area without Ypet signal) to the total mitochondrial area (total TOLLES area) and compared to control conditions (n=4, two-way ANOVA with Dunnett’s correction). Data normalised to control O/A. **E.** Representative images of mitoSRAI POE SH-SY5Y cells pre-treated with DMSO, 10µM NU9056, 100µM MG149 for 3hr and subsequently treated with DMSO or 1µM O/A for 3hr. Cells were immuno-stained for p62 (yellow) and counterstained with DRAQ5 (red) (TOLLES fluorophore in cyan). Scale bar: 20µm. **F.** Quantification of p62 recruitment to the mitochondria in E, calculated as the ratio of mitochondrial to cytoplasmic p62 intensity (ratio of p62 signal intensity inside vs. outside TOLLES), compared to control conditions (n=3, two-way ANOVA with Dunnett’s correction). Data normalised to control O/A. Data shown as mean +/- SD.

During autophagy, mitophagy adapters, such as p62, bind to ubiquitin chains on the OMM, initiating autophagosome formation around the cargo (Johansen, T., *et al*., 2020), ultimately leading to fusion to the lysosomes for degradation (Tanida, I., *et al*., 2008). In order to understand whether the O/A-independant effect of MG149 to induce mitochondrial delivery to the lysosomes is associated with potential increase in autophagosome recruitment to mitochondria, p62 mitochondrial recruitment was next measured. Since the TOLLES fluorophore is localised to the mitochondrial matrix and is resistant to Parkin-dependant degradation and lysosomal degradation, it was used as a p62 mitochondrial co-localisation marker. The O/A-independant effect of MG149 on inducing mitochondrial delivery to the lysosomes (Figures 2A-D) was shown to be associated with an increase in autophagosome recruitment to the mitochondria, as showed by the increase in p62 puncta recruitment to the OMM (Figure 2E,F).

### MG149 treatment stimulates PINK1-dependant mitochondrial clearance

While the PINK1/Parkin-dependant mitophagy process is the best characterized, PINK1/Parkin-independant pathways also play significant roles in mitochondrial clearance (reviewed in Cummins, N., *et al*., 2018 and Whiten, D. R., *et al*., 2021). In order to confirm whether the effects of MG149 on mitophagy were PINK1-dependant or not, a PINK1 CRISPR knockout (KO) POE SH-SY5Y line was generated (PINK1 KO POE). The KO was confirmed by assessment of PINK1 protein levels and PINK1-dependant pUb(Ser65) deposition in PINK1 KO POEs compared to wild type (WT) POEs using immunoblot (Supplementary Figure 2). In line with the crucial importance of PINK1 for O/A-induced mitophagy, no detectable increase in SRAI mitophagy index was observed following treatment of the PINK1 KO POE SHSY5Ys (Figure 3A,B). mitoSRAI PINK1 KO POE cells were pre-treated for 3hr with either 10µM NU9056 or 100µM MG149, prior to treatment with either DMSO or 1µM O/A for 3hr. MG149-treatment elicited no significant change in the SRAI mitophagy index in the PINK1 KO cell line, showing that the mitochondrial clearance initiated by MG149 was dependant on PINK1, both in DMSO and in O/A conditions (Figure 3A,B). To further strengthen this observation, endogenous p62 mitochondrial recruitment was measured in the mitoSRAI PINK1 KO POE cell line, as described above. Treatment with MG149 did not induce any p62 recruitment in the PINK1 KO cells, with or without O/A (Figure 3C,D). Altogther, these data show the MG149-evoked effect on mitochondrial delivery to the lysosomes is PINK1-dependant.

**Figure 3.**
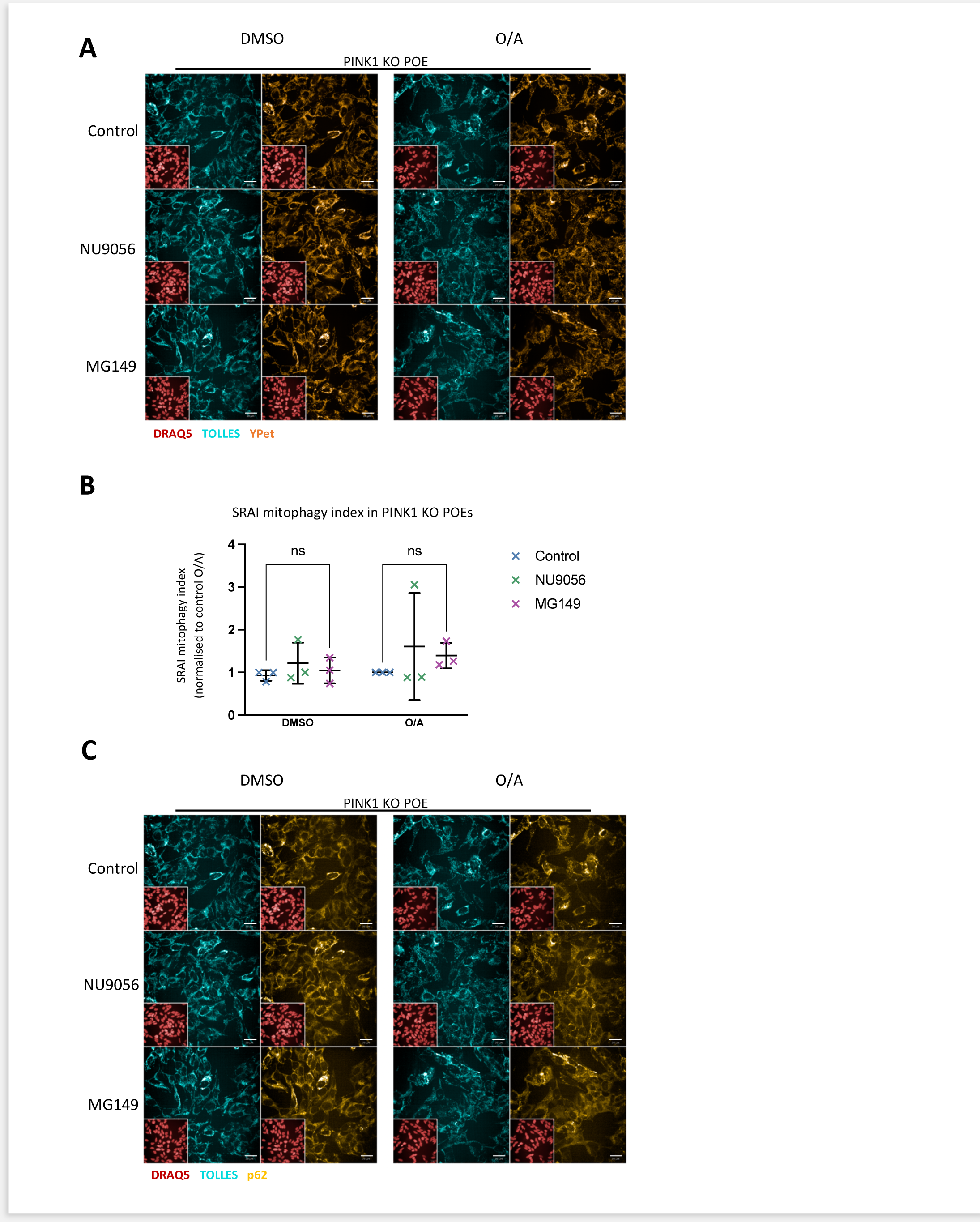
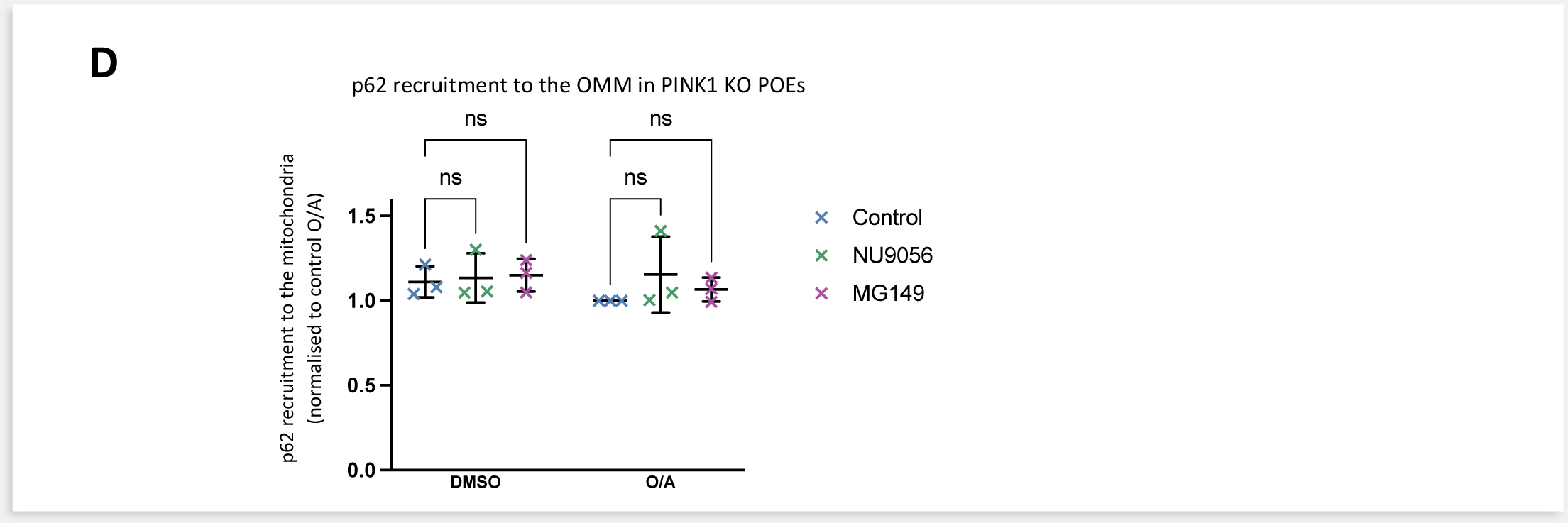
The mitochondrial clearance initiated in MG149 treated cells is dependant on PINK1. **A.** Representative images of PINK1 KO mitoSRAI POE SH-SY5Y cells pre-treated with DMSO, 10μM NU9056, 100μM MG149 for 3hr and subsequently treated with DMSO or 1μM O/A for 3hr. Cells were immuno-stained for p62 (yellow) and nuclei counterstained with DRAQ5 (red) (TOLLES fluorophore in cyan). Scale bar: 20μm. **B.** Quantification of p62 recruitment to the mitochondria in E, calculated as the ratio of mitochondrial to cytoplasmic p62 intensity (ratio of p62 signal intensity inside vs. outside TOLLES), compared to control conditions (n=3, two-way ANOVA with Dunnett’s correction). Data normalised to control O/A. **C.** Representative images of PINK1 KO mitoSRAI POE SH-SY5Y cells pre-treated with DMSO, 10μM NU9056, 100μM MG149 for 3hr and subsequently treated with DMSO or 1μM O/A for 3hr. Cell nuclei were counterstained with DRAQ5 (red). The TOLLES fluorophore (cyan) shows total mitochondria while the Ypet fluorophore (orange) shows the non-lysosomal mitochondria. Scale bar: 20μm. **D.** Quantification of the SRAI mitophagy index in C, calculated as the ratio of the lysosomal mitochondria (TOLLES area without Ypet signal) to the total mitochondrial area (total TOLLES area), and compared to control conditions (n=3, two-way ANOVA with Dunnett’s correction). Data normalised to control O/A. Data shown as mean +/- SD.

### MG149 treatment does not prevent O/A-induced mitochondrial membrane depolarisation

We next attempted to explore possible mechanism(s) of action for MG149-dependant inhibition of PINK1/Parkin-dependant mitophagy initiation. First, we hypothesized that MG149 could inhibit O/A-induced mitochondrial membrane depolarisation required for PINK1 stabilization at the OMM (Yamano, K., *et al*., 2016). In order to test this hypothesis, mitochondrial membrane potential (ΔΨm) was measured in MG149-treated POE SH-SY5Y cells treated with DMSO or O/A, using live confocal fluorescence imaging and TMRM (tetramethyl rhodamine methyl ester) staining (Figure 4A,B). O/A treatment induced mitochondrial depolarisation both in control cells and in MG149 treated cells. In fact, cells pre-treated with MG149 prior to O/A were more depolarised than cells treated with O/A alone, suggesting that the inhibitory effect of MG149 on PINK1-mitophagy initiation was not due a lack of mitochondrial depolarisation. Interestingly, MG149 caused a significant depolarisation of the mitochondrial membrane potential without O/A treatment (Figure 4A, bottom left panel, quantified in Figure 4B). This effect was confirmed looking at OPA1 cleavage in immunoblotting as MG149 alone caused complete cleavage of L-OPA1 into S-OPA1, comparable to observations made following treatment with O/A (Figure 4C,D). The O/A-independant effect of MG149 on ΔΨm is likely to explain the small mitochondrial recruitment and stabilisation of PINK1 at the OMM (Figure 5B,C), the subtle increase in pUb(Ser65) deposition (Figure 1A,B, Supplementary Figure 1B), and ultimately, the increase in mitochondrial delivery to the lysosomes (Figure 2A,B,C,D) following MG149 treatment without O/A.

**Figure 4.**
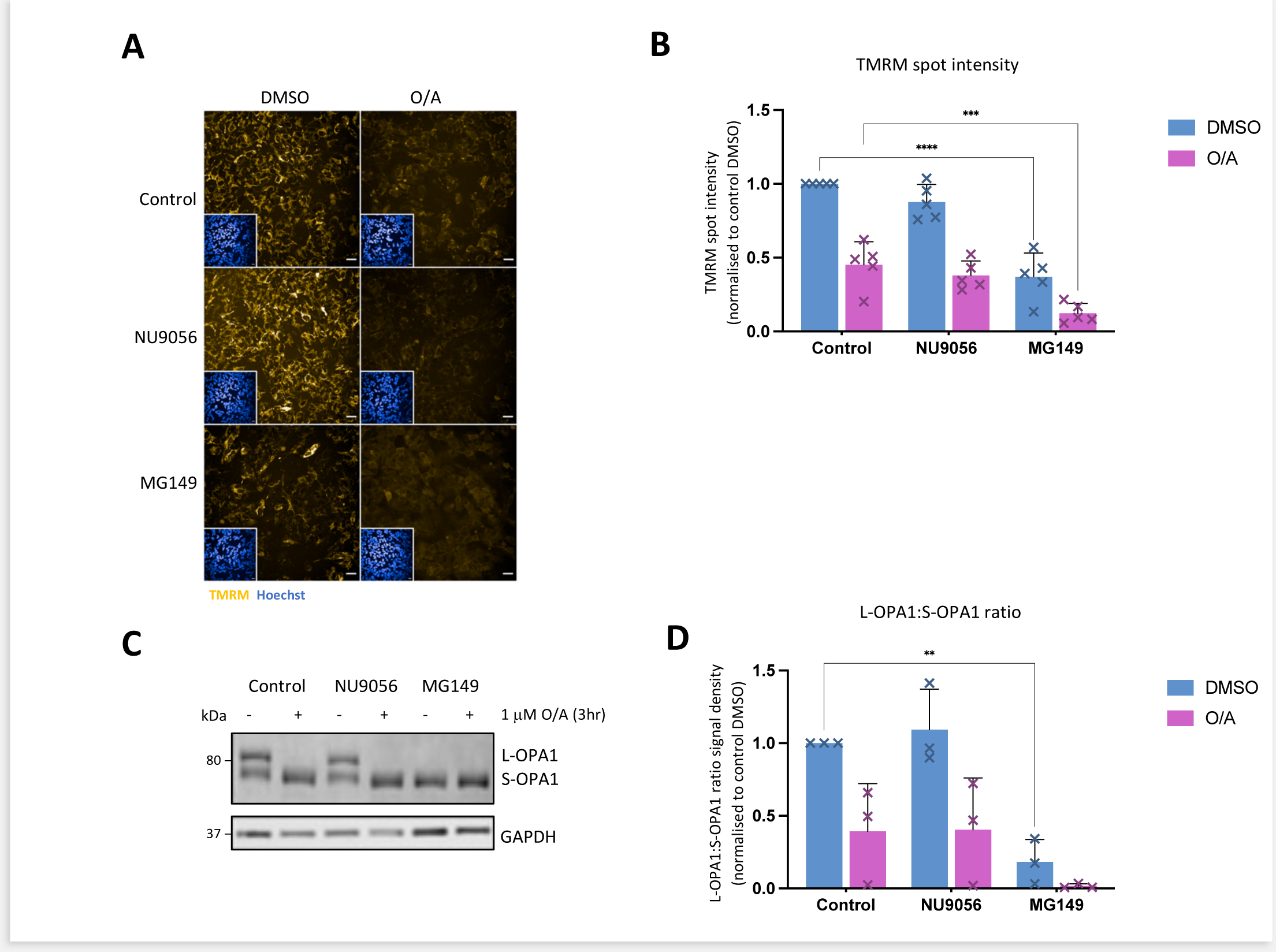
MG149 effectively reduces mitochondrial membrane potential. **A.** Representative images of TMRM (yellow) in POE SH-SY5Y cells pre-treated with DMSO, 10µM NU9056 or 100µM MG149 for 3hr and subsequently treated with DMSO or 1µM O/A for 3hr to induce mitophagy. Nuclei counterstained with Hoechst (blue) Scale bar: 20µm. **B.** Quantification of TMRM spot intensity in A, compared to control conditions (n=5, two-way ANOVA with Dunnett’s correction). Data normalised to control DMSO. **C.** Representative immunoblot of POE SH-SY5Y cells pre-treated with DMSO, 10µM NU9056, 100µM MG149 for 3hr and subsequently treated with DMSO or 1µM O/A for 3hr. Blots were probed for OPA1 and GAPDH. **D.** Quantification of OPA1 signal intensity in C, calculated as the ratio of the signal intensity of L-OPA1 to S-OPA1, and compared to control conditions (n=3, two-way ANOVA with Dunnett’s correction). Data normalized to control DMSO, and GAPDH used as a loading control. Data shown as mean +/- SD.

**Figure 5.**
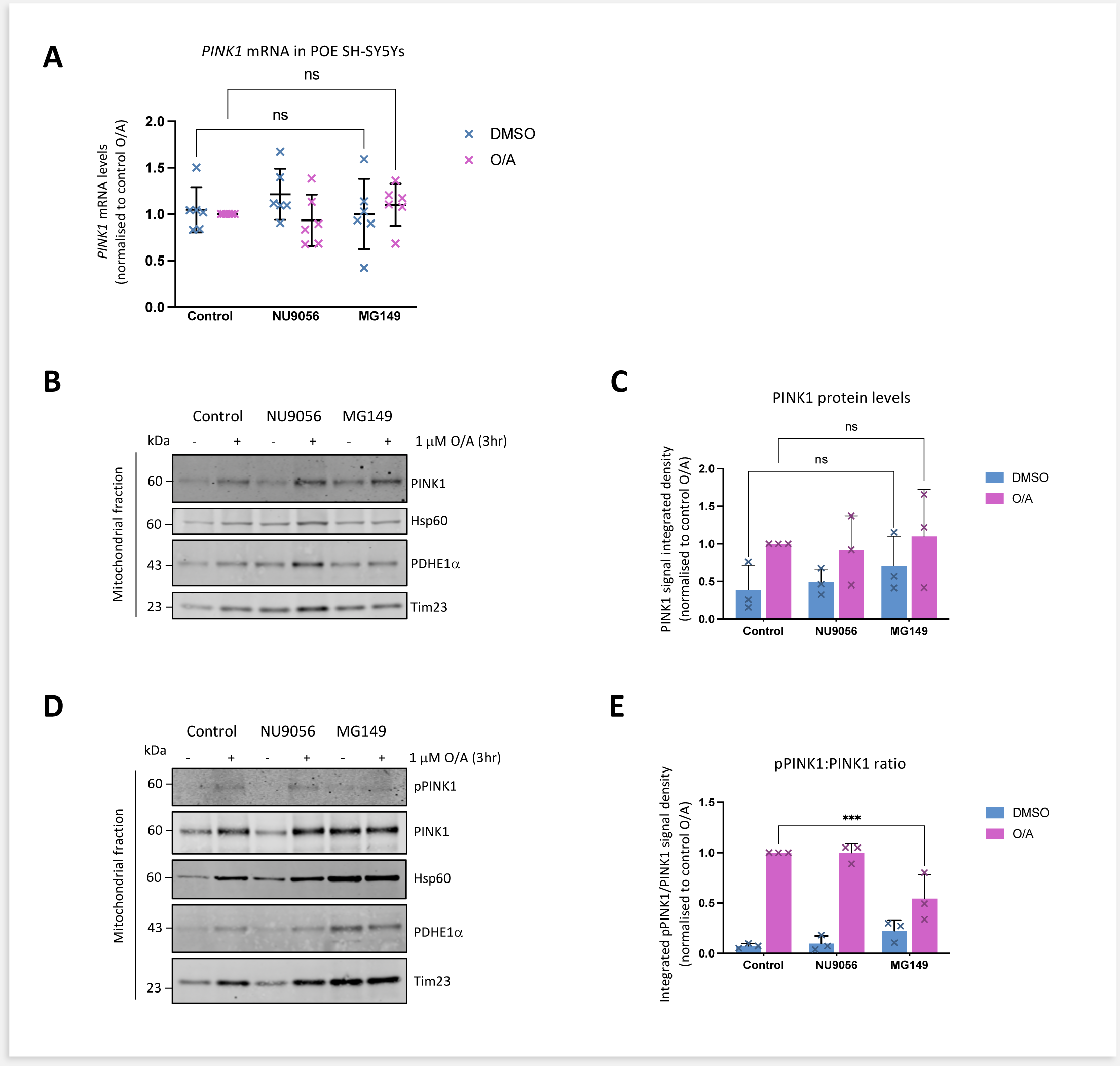
MG149 attenuates PINK1 autophosphorylation. A. Quantification of PINK1 mRNA levels in POE SH-SY5Y cells pre-treated with DMSO, 10µM NU9056, 100µM MG149 for 3hr and subsequently treated with DMSO or 1µM O/A for 3hr, compared to control conditions (n=6, two-way ANOVA with Dunnett’s correction). Data normalised to control O/A. B. Representative immunoblots of mitochondrial fractions from POE SH-SY5Y cells pre-treated with DMSO, 10µM NU9056, 100µM MG149 for 3hr and subsequently treated with DMSO or 1µM O/A for 3hr. Blots were probed for PINK1, Hsp60, PDHE1a, and Tim23. C. Quantification of PINK1 signal intensity in B, compared to controls (n=3, two-way ANOVA with Dunnett’s correction). Data normalised to control O/A, and the PDHE1a used as a loading control. D. Representative immunoblots of mitochondrial fractions from PINK1-HA SH-SY5Y cells pre-treated with DMSO and 100µM MG149 for 3hr and subsequently treated with DMSO or 1µM O/A for 3hr. Blots were probed for pPINK1, PINK1, Hsp60, PDHE1a, and Tim23 E. Quantification of pPINK1 signal intensity in D, calculated as the ratio of pPINK1 to PINK1 signal intensity, compared to controls (n=3, two-way ANOVA with Sídák correction). Data normalized to control O/A and PDHE1a used as a loading control. Data shown as mean +/- SD.

### MG149 treatment does not alter PINK1 gene transcription or PINK1 protein recruitment

We have shown previously that KAT8 siRNA KD POE SH-SY5Ys have reduced levels of *PINK1* mRNA (Soutar, M., *et al*, 2022). Thus, another possible mechanism for decreased mitophagy initiation in MG149-treated cells is a decrease in *PINK1* transcription. To test this hypothesis, we measured the effect of MG149- and NU9056-mediated KAT8/KAT5 and KAT5 inhibition respectively, on *PINK1* transcription, using real-time quantitative polymerase chain reaction (RT-qPCR). Neither of the compound inhibitors significantly impacted *PINK1* mRNA levels (Figure 5A), suggesting that the impairments in PINK1-dependant pUb(Ser65) and pParkin(Ser65) following acute treatment with MG149 were unlikely to be associated with altered transcription of the *PINK1* gene. Next, we assessed whether PINK1 protein accumulation at the mitochondria was altered following treatment of POE SH-SY5Y cells with MG149 using immunoblotting of mitochondrial enriched fractions. In line with the mRNA data, PINK1 protein accumulation was not prevented following MG149 treatment (Figure 5B,C). These data confirm that the decreased levels of PINK1-dependant pUb(Ser65) and pParkin(Ser65) in the MG149-treated samples were not due to a decrease in either *PINK1* mRNA or PINK1 protein levels. In fact, PINK1 protein mitochondrial accumulation was subtly increased following MG149 treatment alone (Figure 5B,C), in line with observations showing MG149 causes mitochondrial depolarisation (Figure 4), subtle deposition of pUb(Ser65) (Figure 1A,B, Supplementary Figure 1B), mitochondrial recruitment of p62 and downstream delivery of mitochondira to the lysosomes in a PINK1-dependant manner (Figure 2, Figure 3), in cells treated with MG149 only.

### MG149 treatment attenuates PINK1 autophosphorylation

Finally, we hypothesized that MG149 treatment could affect PINK1 activation. To test this hypothesis, phosphorylation of PINK1 at serine 228 (pPINK1(Ser228)), an autophosphorylation mark necessary for PINK1 activation, was assessed by immunoblotting mitochondrial enriched fractions from SH-SY5Y cells over-expressing PINK1-HA (PINK1-HA SH-SY5Y) (Okatsu, K., *et al*., 2013). In line with observations in SH-SY5Ys expressing endogenous levels of PINK1 (Figure 5B,C), MG149 alone evoked a subtle increase in mitochondrial PINK1 recruitment in PINK1-HA SH-SY5Ys (Figure 5D). Despite this mitochondrial recruitment of PINK1 in MG149 treated cells, O/A-induced PINK1 phosphorylation at Ser228 was decreased when compared to control, and the mitochondrial PINK1 accumulating with MG149 alone did not appear substantially autophosphorylated either (Figure 5D,E). These data suggest that while MG149 treatment doesn’t prevent mitochondrial depolarisation or PINK1 recruitment, it does reduce PINK1’s autophosphorylation, therefore decreasing its kinase activity and subsequent PINK1-dependant mitophagy initiation.

## DISCUSSION

Dysregulation of the mitochondrial quality control machinery exacerbates toxicity throughout cells, and has been linked to many pathologies, amongst them PD (Brand, M. D., *et al*., 2013). Impairments in mitophagy-associated genes have been identified in several human neurodegenerative diseases such as PD and frontotemporal dementia (FTD) (van der Zee, J., *et al*., 2014), (Lou, G., *et al*, 2020). KAT8 has been identified as both a PD-GWAS candidate risk gene and as being involved in the PINK1/Parkin-dependant mitophagy pathway (Soutar, M., *et al*., 2022). This study explored the impact of MG149, a dual KAT5 and KAT8 inhibitor, on PINK1/Parkin-dependant mitophagy. The absence of a KAT8-specific inhibitor led us to explore the role of MG149 alongside NU9056, a KAT5 specific inhibitor (Li, L., *et al*., 2020).

MG149 significantly decreased all markers of PINK1-dependant mitophagy initiation following O/A treatment (as per pUb(ser65), pParkin(ser65) and MFN2 degradation) without affecting mitochondrial delivery to the lysosomes. Conversely, MG149 also increased PINK1-dependant mitochondrial delivery to the lysosomes independantly of O/A. Together, these data show two opposing but linked consequences of MG149 on PINK1-dependant mitophagy. While MG149 impairs O/A-induced PINK1 autophosphorylation and activation, and subsequent pUb(Ser65) deposition, MG149 alone causes mitochondrial depolarisation, subtle PINK1 accumulation, and subtle pUb(Ser65) deposition. This subtle pUb(Ser65) deposition may be sufficient to recruit autophagy adaptors such as p62 and to promote subsequent delivery of the mitochondria to the lysosomes. In accordance with these findings, past research has linked KAT8 KD to increased autolysosome formation (Hale, C. M., *el al*., 2016). All these effects observed at the mitochondria are nullified in PINK1 KO cell lines, confirming the crucial role of PINK1, and suggesting the KAT8’s activity is intrinsically linked to PINK1.

We have previously shown that KAT8 siRNA KD decreases *PINK1* gene transcription and subsequent PINK1-mitophagy (Soutar, M., *et al*., 2022). Here we show that while MG149 treatment also prevents PINK1-mitophagy initiation, its effect is mediated by altering PINK1 protein kinase activity, rather than *PINK1* gene transcription. Induction of autophagy via starvation was previously associated with KAT8 degradation, subsequent reductions in H4K16 acetylation, and ultimate downregulation of autophagy genes (Füllgrabe, J., *et al*., 2013). The acute, rather than chronic, effect induced by MG149 may explain the lack of transcriptional impact observed. It is possible that prolonged KAT8 compound inhibition may be required for the acetylation mark to be removed and for the *PINK1* gene to be silenced.

All together, our data suggest that KAT8 may play a role in PINK1 mitophagy both by mediating PINK1 gene expression (Soutar, M., *et al*., 2022) and PINK1 protein activity. Our findings may open new avenues for the design of novel therapeutic approaches targeting KAT8 for neurodegenerative diseases such as PD, in which defective PINK1-mitophagy plays a central role.

Playing on the acetylation patterns of histones and proteins may be central in providing new therapies for a variety of diseases. KAT8 inhibition may provide neuroprotective effects for Alzheimer’s disease (AD) by increasing lysosome formation (Yuhong, L., *et al*., 2023).

Interestingly, KAT8 was identified as one of the risk candidate genes for AD (Bellenguez, C., *et al*., 2022). KAT8 inhibition may also have selective toxicity and antiproliferative activity against cancer cells (Zhu, H., *et al*., 2021), (Fiorentino, F., *et al*., 2023) and potentially normalise myocardial functions in the context of cardiovascular diseases and ischemia-reperfusion injury (Yu, L., *et al*., 2018). By looking at MG149 in the context of mitophagy, we have underlined a new area in which modulating KAT8 inhibition may be of use in the study of PD. The lack of specificity of MG149 as a KAT8 inhibitor leaves room for new efficient and potent selective KAT8 inhibitors, such as those described in the recently published study looking for new KAT8 inhibitors for cancer treatment (Fiorentino, F., *et al*., 2023). Hopefully, the development of optimized KAT8 inhibitors may help give a more concrete overview of the therapeutic potential of KAT8 modulation.

## Methods

### Reagents

Oligomycin and antimycin A (inhibitor of mitochondrial complex V and III respectively) were purchased from Sigma-Aldrich (O4876 and A8674 respectively).

For immunoblotting, the following antibodies were used: rabbit anti phospho-ubiquitin (Ser65) (Cell Signaling, 62802, RRID:AB_2799632 1:1000), mouse anti TIM23 (BD Biosciences, 611223, RRID:AB_398755, 1:1000), mouse anti TOM20 (Santa Cruz, sc-11415, RRID:AB_2207533, 1:1000), mouse anti GAPDH (Abcam, ab110305, RRID:AB_10861081, 1:1000), rabbit anti pParkin(Ser65) (Abcam/Michael J.Fox Foundation, MJF17, 1:1000), rabbit anti MFN2 (Cell Signaling, 9482, RRID:AB_2716838 1:1000), mouse anti OPA1 (BD Biosciences, 612607, RRID:AB_399889 1:1000), rabbit anti-LC3B (Novus Biologicals, NB100-2220, RRID:AB_10003146 1:1000) IRDye 680LT donkey anti mouse (LI-COR Biosciences, 925-68022, RRID:AB_2814906, 1:20000), IRDye 800CW donkey anti rabbit (LI-COR Biosciences, 925-32213, RRID:AB_2715510, 1:20000).

Rabbit monoclonal anti PINK1 antibody accessible as described in Soutar, M., *et al*., 2018. For immunofluorescence the following antibodies were used: mouse anti TOM20 (Santa Cruz, sc-17764, RRID:AB_628381 1:1000), rabbit anti phospho-ubiquitin (Ser65) (Cell Signaling, 62802, RRID:AB_2799632 1:1000), mouse anti p62 Lck Ligand (BD Biosciences, 610832, RRID:AB_398151 1:1000), AlexaFluor 488 goat anti rabbit (Invitrogen, A11008, 6 RRID:AB_143165, RRID:AB_ 1:2000), AlexaFluor 568 goat anti mouse (Invitrogen, A11004, 7 RRID:AB_2534072, 1:2000), AlexaFluor 568 anti rabbit (Invitrogen, A11011, RRID:AB_143157, 1:2000), Hoechst 33342 (Thermo Scientific, 62249, 1:10000)), Draq5 (Thermo Scientific, 62254, 1:10000).

### Cell culture

All cells were maintained in culture in a humified 5% CO_2_ incubator at 37°C, in Dulbecco’s modified Eagle medium (DMEM, Gibco, 11995-065) with 4.5 g/L D-glucose and L-glutamine, supplemented with 10% heat-inactivated foetal bovine serum (FBS, Gilbco, 10500-064). Cell medium was refreshed and changed every 3 days.

The human neuronal cell line used for this study was the human neuroblastoma SH-SY5Y cell line: The Flag-Parkin over-expressing (POE) SH-SY5Y cells were a gift from H. Ardley (Ardley H. C., *et al*., 2003), the mitoKeima POE SH-SY5Y cells were a gift from C. Luft (Soutar M., *et al*., 2019), and the PINK1-HA over-expression SH-SY5Y cells were a gift from E. Deas (Deas E., *et al*., 2011 ^1^).

Cells were split at 75% to 90% confluency using 2mL of Trypsin. The flasks were split in such a way that each of their confluency would be approximately the same at the time of manipulations.

### Generation of PINK1 KO POE SH-SY5Ys

PINK1 was functionally knocked out from the POE SH-SY5Y cell line through CRISPR-Cas9 editing by adapting previously published protocols (Cretin, E., *et al*., 2021). Single guide RNA (sgRNA) sequences targeting Exon 1 of *PINK1* (NM_032409) were designed using the Horizon Discovery CRISPR Design Tool (https://horizondiscovery.com/en/ordering-and-calculation-tools/crispr-design-tool). Complementary forward and reverse ssDNA molecules encoding the sgRNA sequence were generated (Sigma Aldrich) inclusive of 5’ overhangs (see Supplementary Table 1 for sequences) suitable for cloning into BbSI digested pSpCas9(BB)- 2A-GFP plasmid (pSpCas9(BB)-2A-GFP Addgene plasmid no. 48138 was a gift from Feng Zhang; RRID:Addgene_48138). A subconfluent 6-well dish of POE SH-SY5Ys were transfected with 1ug of each PINK1_Seq1 and PINK1_Seq2 sgRNA encoding pSpCas9(BB)-2A-GFP plasmid using lipofectamine 3000 (Invitrogen). 24hr post-transfection cells were trypsinised and subjected to fluorescence-activated cell sorting and clonal selection of GFP-positive cells.

Isolated clones were expanded and phenotypically screened for clones which did not show pUb(Ser65) signal after 1.5hr O/A treatment. Briefly, confluent wells of a 96-well, each with a different clone were prepared for immunofluorescence staining of pUb(Ser65) (CST, 62802, RRID:AB_2799632, 1:1000) with IRDye 800CW donkey anti-rabbit secondary (LI-COR Biosciences, 925-32213, RRID:AB_2715510, 1:2000). gDNA was extracted from a positive clone using the Wizard Genomic DNA Purification Kit (Promega) and subjected to whole genome sequencing (Novogene) which revealed homozygous 22bp deletion and frameshift in exon 1 of PINK1 (c.156_177delGGGCGCGGAGCCTCGCAGGGTC, p.A54Sfs*46).

### Generation of mitoSRAI POE SH-SY5Ys

POE SH-SY5Ys stably expressing the mitoSRAI mitophagy sensor (Katayama, H., *et al*., 2020) were established through lentiviral transduction by adapting recently published protocols (Soutar, M. P. M., *et al*., 2022). MitoSRAI cDNA was first PCR amplified from mitoSRAI_pcDNA3 (provided by the RIKEN BRC through the National BioResource Project of the MEXT, Japan; cat. RDB18223) with inclusion of 5’ EcoRI and 3’ NotI restriction sites. The PCR product was then cloned into pLVX-EF1α-IRES-Puro (Clontech, Takara Bio, 631988) using EcoRI and NotI restriction sites to generate pLVX-EF1α-mitoSRAI-IRES-Puro. Lenti-X 293T HEK cells cultured in DMEM 10% FBS media were transfected with pMD2.G, pCMVR8.74 and pLVX-EF1α-mitoSRAI-IRES-Puro at a 1:1:2 molar mass ratio using Lipofectamine 3000 (Invitrogen). The next day, a full media change was performed using DMEM 10% FBS and cells cultured for further 24 h. The lentivirus containing media was collected and diluted 1:2 with DMEM 10% FBS before filtering through 0.44 µm PES filters. pMD2.G (Addgene plasmid no. 12259, RRID:Addgene_12259) and pCMVR8.74 (Addgene plasmid #22036, RRID:Addgene_22036) were gifts from Didier Trono. 1×10^6 POE SHSY5Ys were reverse transduced with the 1:2 lentivirus supernatant dilution in the presence of 10 µg/ml polybrene (Sigma Aldrich). Bulk populations of cells stably expressing mitoSRAI were established through selection with 1ug/ml puromycin (M P BIOMEDICALS UK). Puromycin was maintained during routine culture but withdrawn when seeding for experimental assays.

### Mitophagy Initiation

In order to initiate mitophagy, cells were treated with 1µM oligomycin/ 1µM antimycin for 3hr before collecting / fixing for downstream experiments. Dimethyl Sulfoxide (DMSO, Santa Cruz, sc-202581) was used as a control.

### Small molecule modulators

MG149 targets KAT5 and KAT8 with IC_50_ values of 74µM and 47µM respectively (Li L., *et al*., 2020). NU9056 targets KAT5 with an IC_50_ value of 2µM. Both small molecules were reconstituted in DMSO (Santa Cruz, sc-202581) before treatments.

For immunofluorescence, both compounds were diluted in DMEM (Gibco, 11995-065) so that 25uL of each diluted compound was added per well of a 96-well plate (PhenoPlate 96-well microplate, PerkinElmer, 6055302) to make up desired concentrations.

For immunoblotting, concentrations of 10µM for NU9056 and 100µM for MG149 were added to corresponding wells.

Equal volumes of DMSO (Santa Cruz, sc-202581) were used as no treatment controls.

### Mitochondrial enrichment

Whole cell lysates of POE SH-SY5Y cells were fractionated into cytoplasmic and mitochondria-enriched preparations to facilitate detection of mitochondrial localized proteins of interest. Cells were incubated overnight at −80 °C in mitochondrial homogenization buffer (250mM sucrose, 1mM EDTA, 10mM Tris (pH 7.4) supplemented with protease (cComplete^TM^, EDTA-free Protease Inhibitor Cocktail, Roche, 05056489001) and phosphatase (PhosSTOP^TM^, Roche, 04906837001) inhibitors) before being thawed on ice. The content of the wells were scraped into 1.5ml centrifuged tubes, triturated 20 times and frozen at −80 °C for 1hr. After being thawed on ice, cell debris / non-lysed cells were removed by centrifugation (1500g for 20 mins at 4°C) and the supernatant centrifuged again to pellet the mitochondria (12500g for 20 mins at 4°C). The resulting supernatant was kept, making up the cytoplasmic fraction, and the pellet washed twice with mitochondrial homogenization buffer to remove any cytoplasmic contamination, making up the mitochondrial fraction. Isolated mitochondria pellets were suspended in 1X LDS (NuPAGE™ LDS Sample Buffer (4X), Invitrogen, NP0008) supplemented with 10 mM dithiothreitol (DTT), and sonicated. All LDS preps were heated at 70°C for 10 mins.

### Cell lysis

Cells were lysed in whole-cell lysis buffer (50mM Tris (pH 7.4), 0.1mM EGTA, 1mM EDTA, 0.27M Sucrose, 1% triton-x-100, supplemented with protease (cComplete^TM^, EDTA-free Protease Inhibitor Cocktail, Roche, 05056489001) and phosphatase (PhosSTOP^TM^, Roche, 04906837001) inhibitors) on ice and incubated overnight at −80°C. Lysates were cleared by centrifugation (16000g for 20 mins at 4°C) and protein quantified by BSA assay. LDS preps were made with 10mM DTT prior to heating at 70 °C for 10 mins.

### Western blotting

LDS preparations were separated by SDS-PAGE and transferred to nitrocellulose membrane. Membranes were incubated overnight at 4°C in primary antibodies diluted in 3% milk in PBS-Tween. The following day, membranes were washed and incubated at room temperature for 1hr in secondary antibodies diluted in PBS-Tween. Protein bands were detected using Odyssey CLx LI-COR.

### Immunofluorescence

Cells were fixed with 4% Paraformaldehyde (PFA, Sigma-Aldrich) for 20 mins at room temperature (RT), then blocked and permeabilized with a mix of 10% FBS, 0.25% Triton X-100 in PBS for 1 hr. Cells were then immunostained with primary antibodies diluted in 10% FBS for 2hr at RT. The wells were washed three times using PBS 1X before the secondary antibodies were added and incubated for 1hr: AlexaFluor 568 anti-mouse, 488 anti-rabbit, 568 anti-rabbit secondary antibodies, Hoechst 33342 (Thermo Scientific), and DRAQ5 were all at a 1:2000 dilution in 10% FBS/PBS. The wells were washed three times using PBS 1X, and the plates were imaged using Opera Phenix (PerkinElmer), using the 63X water objective. Images were subsequently analysed using Columbus 2.8 analysis system (PerkinElmer) to measure either the integrated density of pUb(Ser65) within the whole cell (corresponding to the corrected pUb(Ser65) spot intensity x the total spot area), or the mitochondrial to cytoplasmic p62 intensity ratio.

### Measurement of mitochondrial membrane potential using TMRM

POE SH-SY5Y cells were seeded in a 96-well plate (PhenoPlate 96-well microplate, PerkinElmer, 6055302) and treated according to desired conditions (MG149, NU9056, O/A as described above. Before imaging, cells were incubated in 25 nM of Tetramethylrhodamine, Methyl Ester, Perchlorate (TMRM, Sigma Aldrich, T5428) diluted in cell media for 40 mins at 37 °C and 5% CO_2_ (Hoechst 33342 (1:10000) also added to stain the nuclei).

After 40 mins, live cells were immediately imaged on the Opera Phenix (PerkinElmer) at 37 °C and 5% CO_2_, acquiring confocal z-stacks for multiple fields of view across 3 individual wells per experimental condition using the 40X water objective.

Images were then analysed using the Columbus 2.8 analysis system (PerkinElmer) to measure the TMRM spot intensity within the whole cell.

### Measurement of mitochondrial delivery to lysosomes using mitoKeima reporter

mitoKeima expressing POE SH-SY5Y cells were seeded in a 96-well plate (PhenoPlate 96-well microplate, PerkinElmer, 6055302) and treated according to desired conditions (MG149, NU9056, O/A as described above). Before imaging, cell media was replaced with 10% FBS (Gilbco, 10500-064) phenol-free DMEM (Gibco, 31053-028) containing the previously determined concentrations of treatments, and Hoechst 33342 (1:10000) to stain the nuclei. Live cells were immediately imaged on the Opera Phenix (PerkinElmer) at 37 °C with 5% CO_2_, acquiring multiple single-plane fields of view, using the 63X water objective.

Images were then analysed using the Columbus 2.8 analysis system (PerkinElmer) to measure the Keima mitophagy index, which is the ratio between the total area of lysosomal mitochondria (561nm) to the total area of mitoKeima (sum of cytoplasmic (488nm) and lysosomal (561nm) mitoKeima area) per well.

### Measurement of mitochondrial delivery to lysosomes using mitoSRAI

mitoSRAI POE SH-SY5Y cells were seeded in a 96-well plate (PhenoPlate 96-well microplate, PerkinElmer, 6055302), pre-treated with MG149 or NU9056 for 3hr according to desired concentrations, and subsequently with DMSO (Santa Cruz, sc-202581) or O/A for 3hr. Equal volumes of DMSO (Santa Cruz, sc-202581) were used as no treatment controls. After fixed and stained with DRAQ5 to stain the nucleus, as described above, cells were imaged on the Opera Phenix (PerkinElmer).

Images were then analysed using the Columbus 2.8 analysis system (PerkinElmer) to measure the SRAI mitophagy index, which is the ratio between the area of lysosomal mitochondria (corresponding to the total TOLLES area minus YPet signal) and the total mitochondrial area (corresponding to total TOLLES area) per well.

#### Statistical Analysis

Intensity measurements from imaging experiments were normalised as stated for each experiment and presented as a fraction. N numbers are shown in figure legends and refer to the number of independant, replicate, biological experiments. A minimum of 3 biological replicates were used for each experiment. GraphPad Prism 6 (La Jolla, California, USA, RRID:SCR_002798) was used for statistical analyses and graph production. Data were subjected to two-way ANOVA, unless otherwise stated. Data was expressed as the mean ± standard deviation (SD) from replicate experiments and statistical significance was set at p-value of <0.05.

The above methods are detailed in protocols.io: https://www.protocols.io/view/the-impact-of-mg149-a-kat8-inhibitor-on-mitophagy-cuw7wxhn

---

**Supplementary Figure 1.**
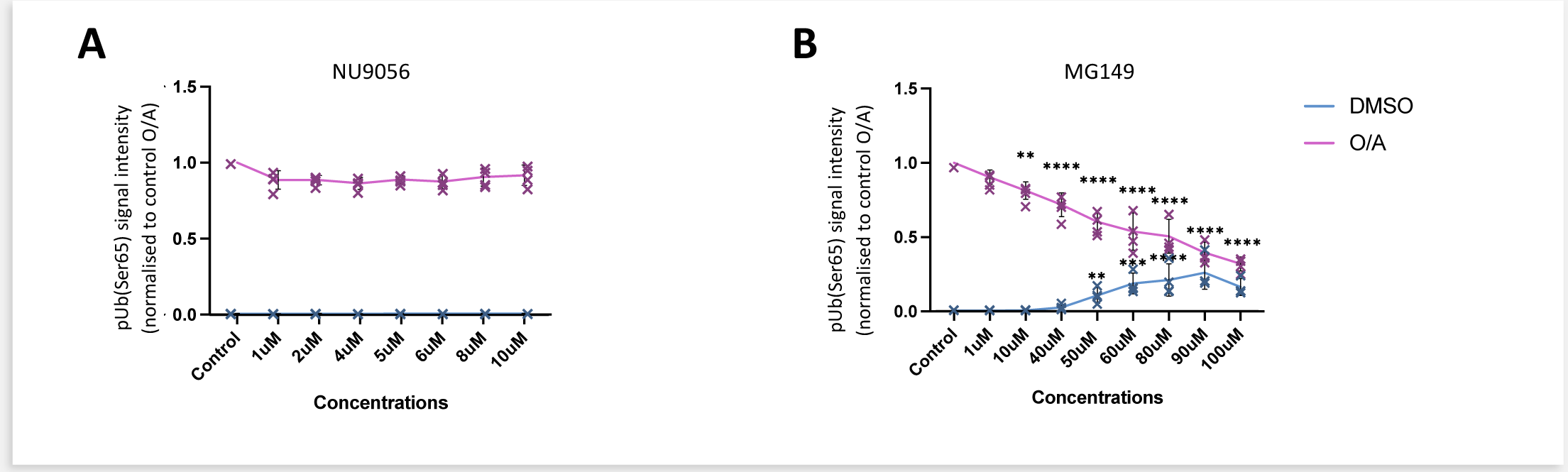
Dose-response of NU9056 and MG149 respectively. **A.** Quantification of NU9056 dose-response looking at integrated pUb(Ser65) signal intensity, compared to control conditions (n=3, two-way ANOVA with Dunnett’s correction). POE SH-SY5Y cells pre-treated with DMSO, 1μM, 2μM, 4μM, 5μM, 6μM, 8μM, 10μM NU9056 or 1μM, 10μM, 40μM, 50μM, 60μM, 80μM, 90μM, 100μM MG149 for 3hr and subsequently treated with DMSO or 1μM O/A for 3hr. Cells were then immunostained with Hoechst and anti-pUb(Ser65). **B.** Quantification of NU9056 dose-response looking at integrated pUb(Ser65) signal intensity, compared to control conditions (n=3, two-way ANOVA with Dunnett’s correction). POE SH-SY5Y cells pre-treated with DMSO, 1μM, 2μM, 4μM, 5μM, 6μM, 8μM, 10μM NU9056 or 1μM, 10μM, 40μM, 50μM, 60μM, 80μM, 90μM, 100μM MG149 for 3hr and subsequently treated with DMSO or 1μM O/A for 3hr. Cells were then immunostained with Hoechst and anti-pUb(Ser65). Data shown as mean +/- SD and significance within DMSO vs O/A treatment groups indicated by *.

**Supplementary Figure 2.**
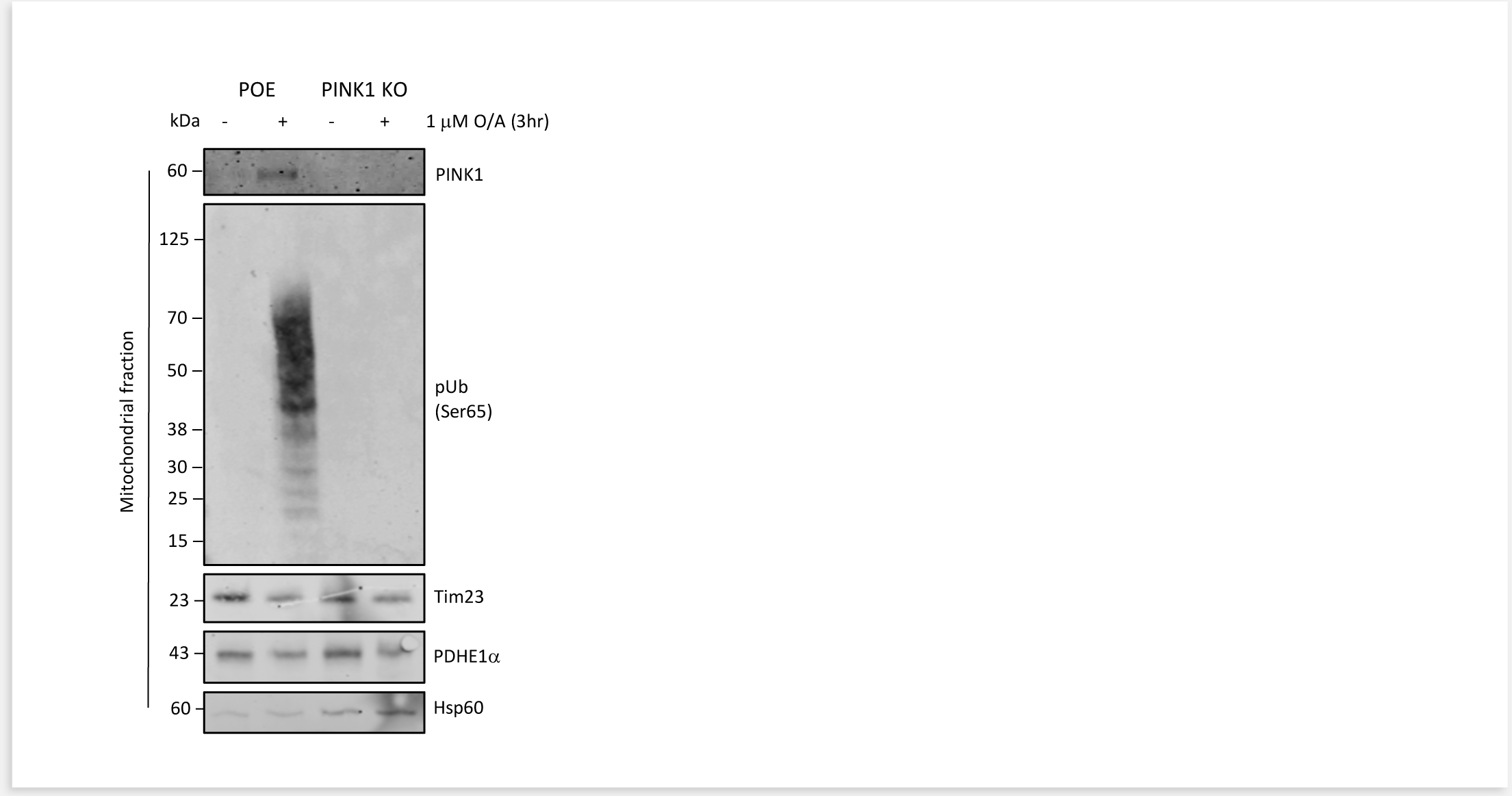
Characterization of PINK1 KO by western blot. Representative immunoblots of mitochondrial fractions from POE SH-SY5Y and PINK1-KO POE SH-SY5Y cells treated with DMSO or 1μM O/A for 3hr. Blots were probed for PINK1, pUb(Ser65), Tim23, PDHE1a, and Hsp60.

**Supplementary Table 1.**
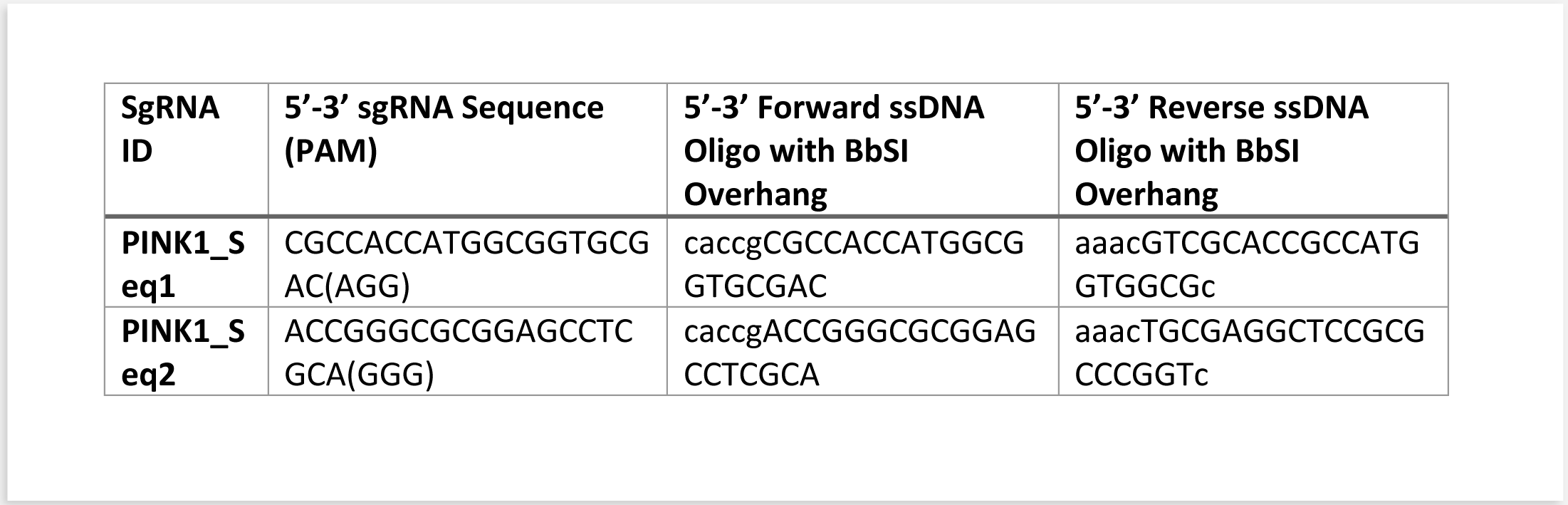
5’-3’ Forward and Reverse ssDNA sequences.

## Notes

https://www.protocols.io/view/the-impact-of-mg149-a-kat8-inhibitor-on-mitophagy-cuw7wxhn

### Competing Interest Statement

The authors have declared no competing interest.

### Summary of Updates

This version of the manuscript has been revised to incorporate insightful comments made by the reviewers after first submission. We have significantly improved the quality and readability of the paper.

